# PREDICTING SAVINGS IN THE METABOLIC COST OF RUNNING WITH AN EXOTENDON

**DOI:** 10.1101/2025.04.15.648814

**Authors:** Jon P. Stingel, Nicholas Bianco, Carmichael F. Ong, Jennifer L. Hicks, Scott L. Delp

## Abstract

The exotendon is a passive device that reduces the energetic cost of running at 2.7 m/s, but its potential benefits at higher speeds remain unknown. Experimental testing is challenging because of the wide range of conditions that must be tested. Here, we use muscle-driven simulations to overcome this challenge and inform exotendon design. We validated a simulation framework that estimates changes in energy expenditure, body kinematics, and muscle activations when simulated subjects run with and without an exotendon. Simulations of people running at 4 m/s with the exotendon that saved energy at 2.7 m/s predicted a 10% reduction in energy cost compared to natural running. We then performed simulations of 25 designs and found that many of the designs saved energy. A longer, stiffer exotendon yielded slightly greater energy savings (12%). Longer and more compliant exotendons offered little savings. We plan to test a limited set of our simulation predictions in an experiment to evaluate their accuracy and assess how an exotendon impacts running performance at 4 m/s. The purpose of this paper is to present the simulation results and to make predictions about the performance of the runner-exotendon system in experiments. This paper has been posted before the experiments have begun to avoid informing the predictions from the experimental results.

## INTRODUCTION

Assistive devices are increasingly used to enhance human motion, but complex human-device interactions make assessing the benefits of each device and its various parameters difficult [1–3]. Additionally, tuning even a few parameters of an assistive device may require an impractical number of physically demanding experimental tests [4]. Previously, an exotendon, a passive spring connecting a user’s shoes, was shown to reduce energy expenditure during slower running (2.7 m/s) [5, 6]. It is unclear if benefits extend to higher speeds, and extensive experimental testing at these speeds is impractical due to physical demands of running fast. Musculoskeletal simulation enables evaluating device parameters without physical testing. In this study, we used musculoskeletal simulations to assess the energetic impact of the exotendon at a faster speed (4 m/s) for a range of exotendon stiffness and lengths..

## METHODS

We developed a planar, 20-degree-of-freedom musculoskeletal model with 18 musculotendon units spanning the lower limbs, four contact spheres per foot, and a spring-damper at the metatarsophalangeal joint [7–11]. To ensure sufficient moment generation capacity for faster running with a reduced muscle set, we doubled the default maximum isometric force values of the Hill-type muscles in the model [11]. We simulated exotendon-assisted running by adding a linear spring between the model’s feet, parameterized by stiffness and slack length. To assess whether optimizing these parameters could improve energetic efficiency at 4 m/s, we tested five stiffness and lengths and evaluated energy expenditure across 25 exotendon parameter combinations.

We generated simulations of natural and exotendon-assisted running at 4 m/s using direct collocation in OpenSim Moco [12]. OpenSim generated muscle-driven kinematic- and kinetic-tracking simulations, with light tracking of joint coordinate trajectories and ground reaction forces while constraining the solution to the target running speed and stride duration. The simulation cost function was formulated as:

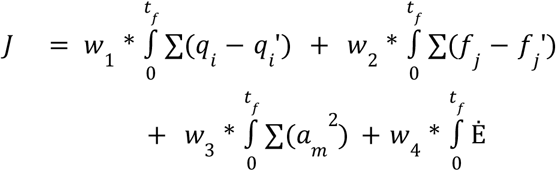

The first term computes kinematic tracking as the difference between the joint coordinate values and speeds for the reference (*q*) and simulated (*q*’) data, summed across all joints, and integrated over the full simulated step duration. The second term computes ground contact tracking similarly between the reference (*f*) and simulated (*f*’) ground contact forces. We minimized the sum of activations squared of all muscles (*a*_*m*_) in the model, integrated over the duration of the step as a third term, as well as the integral of the rate of metabolic energy expenditure (Ė) as the fourth term [13]. Each term in the cost function also has an individual weight (*w*).

As reference data, we used kinematics and kinetics from a published dataset of natural running at 4.0 m/s [14]. We simulated stride durations of 88, 94, 100, and 106% of the duration in the reference data, since the exotendon previously led to changes in stride duration [5]. Results are evaluated at the stride duration with the minimum energetic rate, indicating the model’s optimal performance for that set of device parameters.

## RESULTS AND DISCUSSION

The simulation predicted that the exotendon that saved energy at lower speed (2.7m/s) would also save energy runners at 4 m/s. We found a 10% (0.73 J; 1.13 W/kg) reduction in rate of energy expenditure compared to natural running (Table 1), a larger change than the 6.4% savings reported at 2.7 m/s experimentally [5]. The savings primarily occurred during stance, with the quadriceps muscles having the largest savings during stance (1.53 W/kg). Simulations showed that most of the exotendon parameter combinations we simulated would be beneficial for runners at 4 m/s (Table 1). We saw that exotendons that had a high stiffness and short slack length increased the energetic burden, while long exotendons with low stiffness had little effect. The exotendon combination with the largest savings was one with a slack length of 0.4305 cm and a stiffness of 240 N/m. The exotendons with the five largest reductions in energy expenditure had an average peak tension of 102±17 N, though a majority of the exotendon parameter combinations were beneficial compared to natural running (Fig. 1). These results suggest that there is some flexibility in the amount of force that a runner can benefit from when using the exotendon, but that there may be a more optimal range of force input from the greatest energetic savings.

**Table 1.** Percent changes in average energy expenditure for each exotendon parameter combination at 4.0 m/s compared to natural running. Increases (+) are highlighted in pink, decreases (-) in green. Stiffness values (k, rows) range from 25-200% of the original 120 N/m, and lengths (l, columns) from 25-200% of the original 0.29 m. The original parameters (l = 0.29 m, k = 120 N/m) are outlined in black.

| % change in energy expenditure Natural to Exotendon 4.0 m/s |  | Exotendon length (m) |  |  |  |  |
| --- | --- | --- | --- | --- | --- | --- |
|  |  | l=0.07 | l=0.14 | l=0.29 | l=0.43 | l=0.57 |
| Exotendon stiffness (N/m) | k= 30 | -3.9 | -3.2 | -2.4 | -0.7 | -0.1 |
|  | k= 60 | -9.9 | -9.1 | -6.2 | -3.9 | -1.9 |
|  | k= 120 | -11.4 | -11.7 | -10.0 | -9.6 | -5.4 |
|  | k= 180 | -9.0 | -10.8 | -12.1 | -11.7 | -8.8 |
|  | k= 240 | 1.3 | -4.8 | -11.7 | -12.2 | -11.1 |

**Fig 1:**
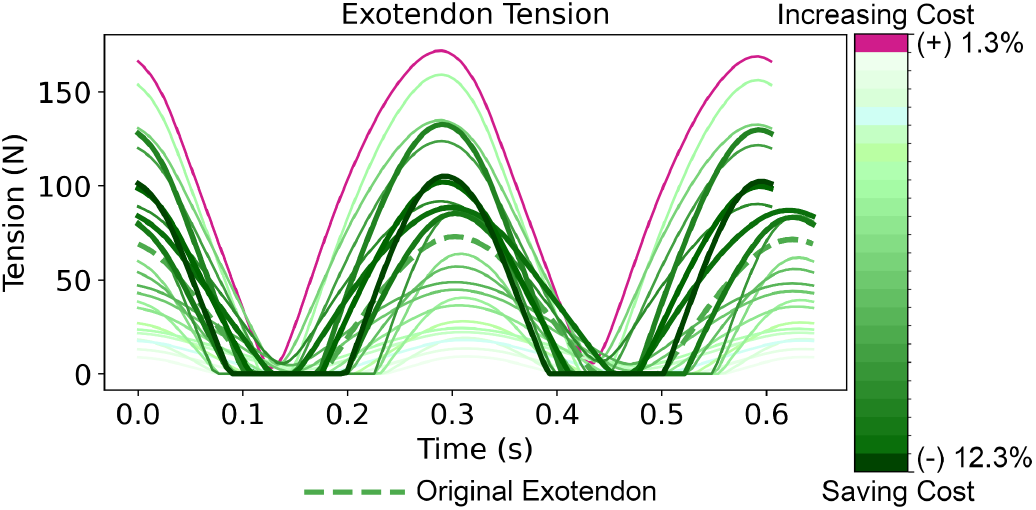
Exotendon tension force over a stride time is shown, with color indicating energetic impact: darker green for savings, darker pink for increased expenditure, and lighter colors for minimal change. The original exotendon is marked with a dashed line.

We plan to test these predictions in an upcoming experiment. In the experiment, runners will complete speed controlled trials at 4 m/s on a treadmill, to assess their energetic changes after training with each of the devices. In the experiment, we plan to test the following exotendons shown in Table 1:

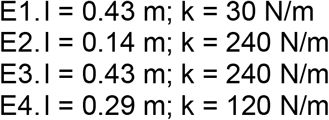

Exotendon E1 represents a longer, compliant exotendon that will supply only a small amount of force. E2 represents a shorter, stiffer exotendon that would supply a much higher force to the runner. E3 signifies the optimal exotendon from the simulations, and E4 is the exotendon that represents the original set of design parameters.

We hypothesize that the optimal and original tendon will save energy compared to natural. If correct, this shows that there are multiple effective exotendon configurations, and the original exotendon is beneficial at a range of running speeds. We hypothesize that the optimal and original are more effective than the short and stiff and long and compliant exotendons. This would show that the simulations can identify exotendon parameters that are energetically beneficial.

We will also evaluate how an exotendon might affect the runners’ performance in a 5 kilometer run. Runners will complete an unassisted, maximal-effort 5K run on a standard outdoor track at the beginning of the experiment. They will then complete the same run while using the exotendon that supplied them the greatest energetic benefit. We will compare their run times.

We hypothesize that the exotendon with the maximum savings will significantly improve 5K race times on the track.

## CONCLUSIONS

Our musculoskeletal simulations suggest that an exotendon can reduce energetic burden at 4 m/s, extending its benefits to faster running speeds. The results identified several effective exotendon parameters at this speed. This simulation framework enables efficient testing and optimization of device parameters, which can guide future experimental studies and reduce physical burden. We plan to test the exotendon in a prospective experiment at 4 m/s to evaluate its efficacy and our model predictions.

## REFERENCES

1. Beck ON et al., R Soc Open Sci, 9(1):211799, 2022.

2. Greenemeier LIn: Sci. Am. Accessed 29 Apr 2024

3. Beck ON et al., Sci Rep, 10(1):17154, 2020.

4. Poggensee KL et al., Sci Robot, 6(58):eabf1078, 2021.

5. Simpson CS et al., J Exp Biol, jeb.202895, 2019.

6. Stingel JP et al., IEEE Robot Autom Lett, 8(10):6267–6274, 2023.

7. Pariser KM et al., J. Biomech. Eng. 144:

8. Ong CF et al., PLOS Comput Biol, 15(10):e1006993, 2019.

9. Bianco NA et al., PLOS ONE, 17(1):e0261318, 2022.

10. Rajagopal A et al., IEEE Trans Biomed Eng, 63(10):2068–2079, 2016.

11. Haralabidis N et al., PeerJ, 9:e10975, 2021.

12. Dembia CL et al., PLOS Comput Biol, 16(12):e1008493, 2020.

13. Bhargava LJ et al., J Biomech, 37(1):81–88, 2004.

14. Hamner SR et al., J Biomech, 46(4):780–787, 2013.

